# Genetic dissection of cyclic pyranopterin monophosphate biosynthesis in plant mitochondria

**DOI:** 10.1101/234450

**Authors:** Inga Kruse, Andrew Maclean, Lionel Hill, Janneke Balk

## Abstract

Mitochondria play a key role in the biosynthesis of two metal cofactors, iron-sulfur (FeS) clusters and molybdenum cofactor (Moco). The two pathways intersect at several points, but a scarcity of mutants has hindered studies to better understand these links. We screened a collection of sirtinol-resistant *Arabidopsis thaliana* mutants for lines with decreased activities of cytosolic FeS enzymes and Moco enzymes. We identified a new mutant allele of *ATM3*, encoding the ATP-binding cassette Transporter of the Mitochondria 3 (systematic name ABCB25), confirming the previously reported role of ATM3 in both FeS cluster and Moco biosynthesis. We also identified a mutant allele in *CNX2, Cofactor of Nitrate reductase and Xanthine dehydrogenase 2*, encoding GTP 3′,8-cyclase, the first step in Moco biosynthesis which is localized in the mitochondria. A single nucleotide polymorphism in *cnx2-2* leads to substitution of Arg88 with Gln in the N-terminal FeS cluster-binding motif. *cnx2-2* plants are small and chlorotic, with severely decreased Moco enzyme activities, but they performed better than a *cnx2-1* knockout mutant, which could only survive with ammonia as nitrogen source. Measurement of cyclic pyranopterin monophosphate (cPMP) levels by LC-MS/MS showed that this Moco intermediate was below the limit of detection in both *cnx2-1* and *cnx2-2*, and accumulated more than 10-fold in seedlings mutated in the downstream gene *CNX5.* Interestingly, *atm3-1* mutants had less cPMP than wild type, correlating with previous reports of a similar decrease in nitrate reductase activity. Taken together, our data functionally characterise *CNX2* and suggest that ATM3 is indirectly required for cPMP synthesis.

## Introduction

Iron-sulfur clusters (FeS) and molybdenum cofactor (Moco) are two covalently bound metal cofactors that mediate different types of redox reactions. Both FeS and Moco must be synthesized *de novo* in cells because they are chemically unstable, particularly under aerobic conditions [1,2]. There are only a few Moco enzymes known in eukaryotes: xanthine dehydrogenase in purine catabolism, aldehyde oxidase, sulfite oxidase, the mitochondrial amidoxime-reducing components mARC1 and mARC2, and nitrate reductase [3,4]. Cells harbour many different FeS enzymes. At least 100 FeS proteins are found in the model plant *Arabidopsis thaliana*, which are involved in respiration, photosynthesis, DNA metabolism and a wide range of metabolic pathways [1,5].

Moco consists of a pyranopterin ring structure with two sulfur atoms forming an enedithiolate group that coordinates molybdate [6]. The biosynthesis of Moco has been primarily characterized in bacteria, the fungus *Aspergillus nidulans*, Arabidopsis and humans [2,7]. The pathway is largely conserved across all domains of life but has been lost from Baker’s yeast. Moco biosynthesis starts with the condensation of GTP into cyclic pyranopterin monophosphate (cPMP) by the consecutive action of GTP 3′,8-cyclase and cPMP synthase [8], encoded by *CNX2* and *CNX3* in plants, respectively (Figure 1). The homologs in bacteria are *MoaA* and *MoaC*, and *MOCS1A* and *MOCS1B* in human. In eukaryotes, the synthesis of cPMP is localized in the mitochondrial matrix whereas the next steps occur in the cytosol [9,10]. Two sulfurs are inserted into the pterin mediated by the concerted action of CNX5, CNX6 and CNX7. The CNX5 (MOCS3) protein first adenylates the C-terminal glycine of CNX7 (MOCS2A) and subsequently replaces the adenylate group with a sulfur, to form a thiocarboxylate. CNX7 binds to CNX6 (MOCS2B) to catalyse sulfur transfer to the third pterin ring, which is repeated for a second sulfur to form enedithiolate. In both plants and humans, CNX5 / MOCS3 also mediates thio-modification of cytosolic tRNA_UUU_ (Lys), tRNA_UUC_ (Glu) and tRNA_UUG_ (Gln), with URM11 (URM1 in humans) acting as sulfur acceptor [11,12]. Finally, CNX1 inserts Mo into the enedithiolate group. One more maturation step is required for a specific class of Moco enzymes, aldehyde oxidases and xanthine dehydrogenases, namely sulfuration of the molybdenum atom by a highly specialized cysteine desulfurase, ABA3 in Arabidopsis and MOCOS in mammals [7,13,14].

**Figure 1.**
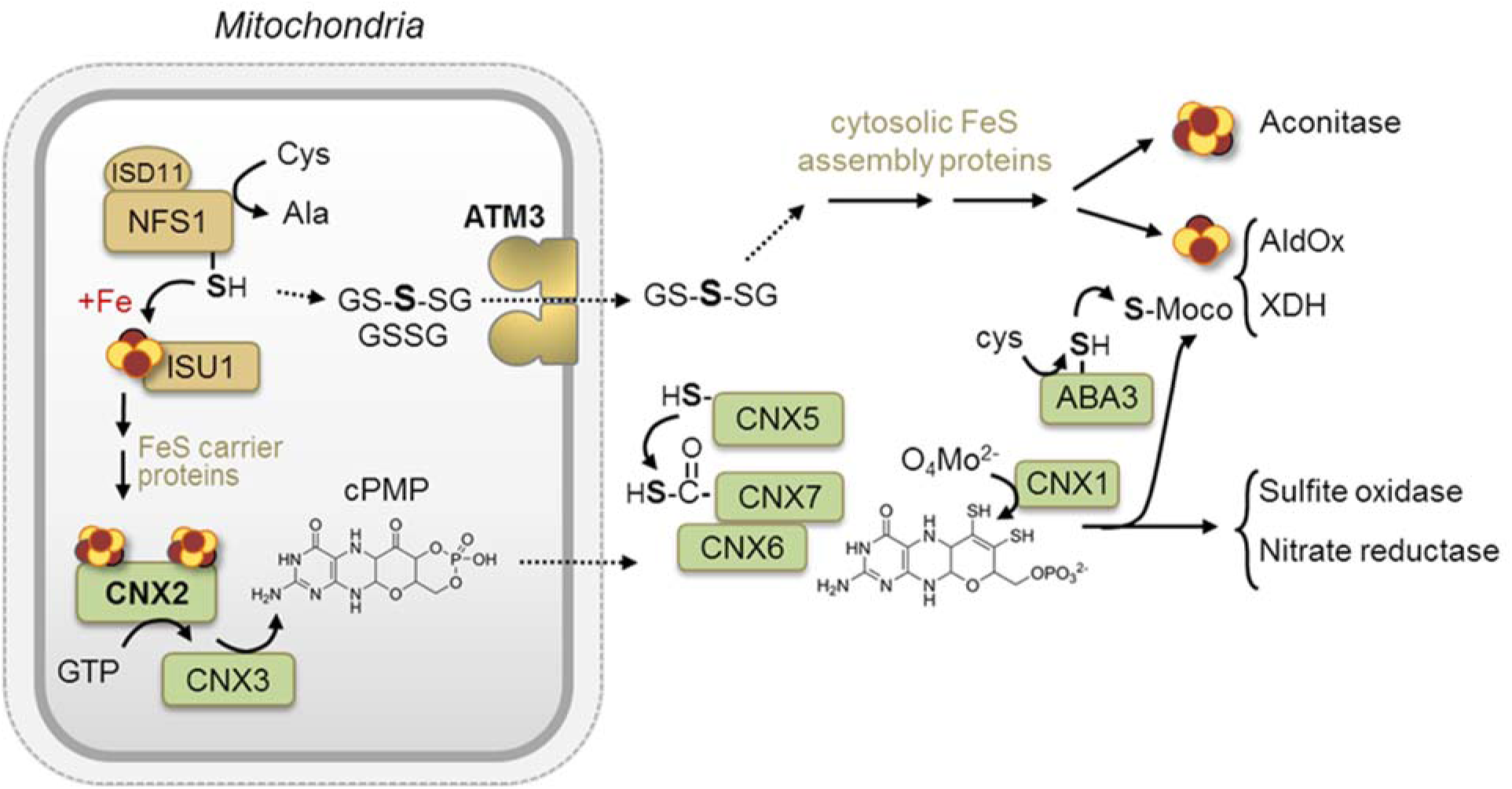
Diagram of the pathways for FeS cluster assembly (light brown) and Moco biosynthesis (green) in Arabidopsis. Solid arrows indicate chemical transition based on independent experimental evidence; dashed arrows indicate more speculative steps. In human cells, cytosolic NFS1 is thought to provide sulphur to the CNX5 homologue. cPMP, cyclic pyranopterin monophosphate; GSSG, glutathione disulphide; GS-S-SG, glutathione trisulfide. For protein acronyms, please see the main text.

FeS clusters occur mostly as rhombic Fe_2_S_2_ or cubane Fe_4_S_4_ clusters. Mitochondria and plastids are the main sites of FeS cluster biosynthesis. Sulfur is provided by a cysteine desulfurase, which is NFS1 in the mitochondria and NFS2 in the plastids. The sulfur is directly transferred from the enzyme active site to the scaffold protein, ISU1 (ISCU) in mitochondria, where it is combined with iron. Additional carrier proteins are required for cluster transfer to target FeS proteins, see [15,16] for reviews. Interestingly, the activities of cytosolic and nuclear FeS enzymes depend on NFS1 in the mitochondria in both plants and humans [1,17]. The plastid-localized NFS2 does not play any role in the maturation of cytosolic FeS enzymes, nor does ABA3 [17], which only serves to sulfurate Moco. A mitochondrial ATP Binding Cassette (ABC) transporter conserved in plants (ATM3/ABCB25), mammals (ABCB7) and yeast (Atm1) is also required for cytosolic FeS enzymes, and has been suggested to export a sulfur-containing compound from the mitochondria to the cytosol. *In-vitro* studies to identify the substrate of ATM3/Atm1 have focussed on glutathione derivatives and showed that they can transport glutathione disulfide and, in the case of Atm1, glutathione trisulfide [18].

The Moco and FeS biosynthetic pathways intersect in several places, as reviewed for bacteria [19] and modified here for Arabidopsis (Figure 1). Firstly, NFS1 provides sulfur for both FeS clusters and Moco in bacteria and in human cells. Studies in HeLa cells showed that a cytosolic form of NFS1 physically interacts with MOCS3 to provide a sulfur atom for subsequent relay to MOCS2A [10,20]. In plants, there is as yet no evidence for a cytosolic pool of NFS1, which appears to be exclusively localized to mitochondria based on GFP studies [21,22] and large-scale proteomics data (www.suba.live). Secondly, CNX2 / MOCS1A is an FeS enzyme depending on two Fe_4_S_4_ clusters, which are assembled by the mitochondrial FeS cluster assembly pathway. Thirdly, ATM3 in Arabidopsis is required for the activity of both Moco and cytoslic FeS enzymes [9,23]. It has been suggested that cPMP is a substrate of ATM3, but this idea has not been tested in transport assays, and it should be noted that cPMP can cross membranes in co-culture experiments [24] or therapeutic treatments for Moco deficiency [25]. A fourth convergence point of the FeS and Moco pathways are enzymes that bind both types of cofactor, the cytosolic enzymes aldehyde oxidase and xanthine dehydrogenase.

To better understand the cross-over points of the FeS and Moco pathways, we screened a sirtinol mutant collection for mutants with decreased aldehyde oxidase activities. We identified and characterised a new *atm3* allele and a viable allele of *cnx2* in Arabidopsis. A new LC-MS/MS method was developed to measure cPMP in plant samples, which revealed that cPMP was strongly decreased in *cnx2* mutants but accumulated in a *cnx5* mutant. cPMP was also decreased in *atm3-1*, providing a biochemical explanation for partially decreased Moco enzyme activities and why *atm3* mutant alleles are found together with mutants in Moco biosynthesis.

## Experimental

### Plant material and genetic analysis

*Arabidopsis thaliana* ecotype Columbia (Col-0) or Landsberg (Ler) were used as wild-type controls. The *atm3-1*, *atm3-2* and *atm3-4* lines in the Col-0 background have been described previously [23]. The *cnx5 (sir1)* mutant has also been reported previously [26], containing substitution of the conserved Ser149 residue to Phe, caused by a C to T point mutation which we confirmed by sequencing. The *xd22* and *xd105* lines were a gift from Florian Bittner and originate from a sirtinol resistance screen carried out on ethyl methanesulfonate (EMS)-mutagenized Arabidopsis (Ler) by Dai *et al.* (2005) [27]. The T-DNA insertion line *cnx2-1* (SALK_037143 in Col-0 background) was obtained from the Nottingham Arabidopsis Stock Centre. Genotyping was carried out using gene-specific primers CNX2 2F and CNX2 2R for the wild-type *CNX2* allele and CNX2 2F with LBb1.3 for the T-DNA insertion. A PCR/restriction assay was designed to detect the *xd22* point mutation, which removes a BsaWI site: the PCR product generated with primers CNX2endog 1F and CNX2endog 1R was digested with BsaWI, resulting in 3 fragments for wild-type *CNX2* and 2 fragments for the *cnx2-2* allele.

### Plant growth

All seeds were vernalised for 2 days at 4°C. Seeds were sown directly onto Levington’s F2 compost, or they were surface sterilised using chlorine gas and spread on ½-strength Murashige and Skoog (MS) medium containing 0.8% (w/v) agar. When indicated 1% (w/v) sucrose was included. To overcome growth impairment due to diminished nitrate reductase activity, seedlings were grown on plates containing 2.5 mM (NH_3_)_2_succinate as the only nitrogen source and buffered at pH 5.6 with 2.5 mM 2-(N-morpholino)ethanesulfonic acid-KOH [28]. Plants were grown under long-day conditions (16 hours light, 8 hours dark) at 22°C and a light intensity of 180-200 μmol m^−2^ s^−1^ for growth on soil or 120-160 μmol m^−2^s^−1^ for growth on agar medium. Plants for isolation of mitochondria were grown under short-day conditions (8 hours light, 16 hours dark) to maximize the amount of leaf material.

### Mapping mutations by whole genome sequencing

Coarse mapping of the *xd22* mutation was performed using simple sequence length polymorphism (SSLP) markers. Whole genome sequencing was carried out by the Earlham Institute (Norwich Research Park) using the Illumina GAIIx platform with 80 bp paired-end reads and ≥ 30x coverage. Sequence assembly and alignment was performed by two independent bioinformatics methods (1) Bowtie [29] hosted on http://bowtie-bio.sourceforge.net/index.shtml and the Integrative Genomics Viewer [30] http://software.broadinstitute.org/software/igv/; (2) Cortex [31] hosted on http://cortexassembler.sourceforge.net/index.html. The reference Ler genome was obtained from http://mus.well.ox.ac.uk/19genomes/. The single nucleotide polymorphisms (SNPs) were filtered for EMS mutant exchanges (G>A and C>T) and a stringent level of homozygosity was applied (>90% of reads supporting the variant). This identified 2065 SNPs genome-wide and 54 within the mapping interval of chromosome 2 (~12799630-16291977).

### RNA extraction and cDNA synthesis

RNA was extracted from plant material using a QIAGEN extraction kit following the manufacturer’s instructions. cDNA synthesis was performed using the SuperScript TM III Reverse Transcriptase (Invitrogen) and an oligo-dT primer as per manufacturer’s instructions. RT-PCR was performed using the primers ACT2 F2 and ACT2 R2 for *ACTIN2*, ATM3 RT-F1 and ATM3 RT-R1 for *ATM3* and CNX2 3F and CNX2 3R for *CNX2*.

### Molecular cloning and plant transformation

For complementation of the *cnx2-2* phenotype, a genomic fragment including the promoter region from position −1224 and UTRs was amplified by PCR using the primers A1F and A1R and Phusion polymerase as per manufacturer’s instructions (New England Biolabs). The PCR fragment was cloned in a pUC-derived cloning plasmid using HindIII and KpnI to cut the vector and an In-Fusion Kit (Takara) to insert the fragment. After confirming the sequence, the gene fragment was cut out using AscI and PacI and ligated into a pBIN-derived binary vector containing a hygromycin resistance marker. *cnx2-2* plants were transformed using the floral-dip method and *Agrobacterium tumefaciens* (strain GV3101). Successfully transformed plants were selected on medium containing 25 μg ml^−1^ hygromycin B.

### Treatments with cPMP

Chemically synthesized cPMP•HBr•2H_2_O [32] was provided by Alexion Pharmaceuticals, CT, USA and dissolved in dimethyl sulfoxide. All solutions were saturated with nitrogen gas to minimize oxidation of cPMP. For application of cPMP to the roots, a 1 mM stock solution of cPMP was diluted 10-fold in ½ MS medium containing 1% (w/v) sucrose, and this was injected into the ½ MS agar plates around the roots of 2-week-old *cnx2-1* seedlings. For vacuum infiltration, a 10 mM cPMP stock solution was diluted 25-fold in 50 mM KPO_4_ pH 7.2, 5 mM ascorbic acid and 0.005% (v/v) Silwet L-77. Four-week-old *cnx2-2* plants were held up-side-down into the solution and vacuum was applied at −30 kPa for 1 min. Transparent patches in the leaves indicated that the solution had entered the intracellular spaces. These patches disappeared after 2 hours and no tissue damage was visible by the naked eye.

### Protein blot analysis

Mitochondria were purified from 4-week-old rosette leaves using differential centrifugation and density gradients [33]. Mitochondrial proteins were solubilized in sample buffer (0.125 M Tris-HCl pH 6.8, 2% (w/v) sodium dodecyl sulfate, 10% (v/v) glycerol, 5% (v/v) 2-mercaptoethanol, 0.1% (w/v) bromophenol blue) and separated by standard sodium dodecyl sulfatepolyacrylamide gel electrophoresis (SDS-PAGE). This was followed by transfer of the proteins to nitrocellulose membrane and immunolabelling with specific antibodies. Antibodies against ATM3 and TOM40 were as previously described [23,34].

### Enzyme assays

All enzyme activities were carried out on 4-week-old rosette leaves. Activity assays for aldehyde oxidase (AldOx) and xanthine dehydrogenase (XDH) were performed using in-gel assays as previously described [23]. Nitrate reductase activity was measured following the production of nitrite, essentially as reported by [35]. Aconitase activities were visualized using an in-gel activity staining method for small tissue samples, as described by Bernard et al. (2009). To confirm equal protein loading, aliquots of the protein extracts were separated by SDS-PAGE and stained with Coomassie, or transferred to nitrocellulose followed by Ponceau-S staining of the membrane.

### cPMP measurement by LC-MS/MS

Sample preparation and conversion of cPMP to compound Z was adapted from [8]. Whole seedlings or rosette leaves were ground with 1 volume ice-cold 10 mM Tris-HCl pH 7.2 per gram fresh weight. Insoluble debris was removed by centrifugation at 16,000 x *g* for 10 min at 4°C. Ninety μl of sample was oxidized by adding 10 μl acidic I_2_/KI (1% / 2% (w/v)) and incubated at room temperature for 20 min to convert cPMP to compound Z. Samples were centrifuged at 16,000 x *g* for 10 min at room temperature to remove precipitated protein. The supernatant was diluted 10-fold with 50% (v/v) acetonitrile and 5 μl was run on a XEVO TQS tandem quadrupole mass spectrometer. Separation was on a 100 x 2.1 mm, 2.6 μm particle size Accucore™ 150 Amide HILIC LC column (Thermo) using the following gradient of 0.1% (v/v) formic acid in H_2_O (Solvent A) versus acetonitrile (Solvent B), run at 500 μl min^−1^ and 40°C: 0 min, 88% B; 7.5 min, 73% B; 14 min, 50% B; 14.5 min, 50% B; 14.6 min, 88% B; 20 min, 88% B. Compound Z formed a hydrogen adduct *m*/*z* 344, and was monitored by the transition 344 to 217.9 at 25V collision energy. Spray chamber conditions were 500°C desorbation temperature, 900 l.h^−1^ desolvation gas, 150 l.h^−1^ nebulizer gas, 7 bar nebulizer pressure, and a capillary voltage of 3.9 kV; the cone voltage was 30 V. The concentration of cPMP was estimated by the method of standard addition, using a series of wild-type samples spiked with increasing amounts of synthetic cPMP.

### Databases and bioinformatic analysis

Amino acid sequences were obtained from the TAIR database, www.arabidopsis.org. Protein sequences were obtained from the UniProt server (www.uniprot.org). Unless otherwise stated, sequence alignments were generated using ClustalW omega (www.ebi.ac.uk/Tools/msa/clustalo). Shading was performed using the Boxshade server www.ch.embnet.org/software/BOX_form.html. Prediction of transmembrane helix formation was performed using TMHMM2 (www.cbs.dtu/services/TMHMM-2.0) [36,37].

### Gene identifiers

ABA3, AT1G16540; ABCB25 / ATM3, AT5G58270; ACTIN2, AT3G18780; CNX1, AT5G20990; CNX2, AT2G31955; CNX3, AT1G01290; CNX5, AT5G55130; CNX6, AT2G43760; CNX7, AT4G10100; TOM40, AT3G20000.

### Statistical analysis

Statistical analysis was performed using Genstat (version 18). Specific statistical information is given in figure legends.

## Results

### Selection of sirtinol-resistant mutants with FeS and Moco defects

In plants, sirtinol has been used to screen for mutants in auxin signalling, an important plant growth hormone [26,27]. Sirtinol undergoes a series of metabolic transformations to 2-hydroxy-1-naphthaldehyde, which is then oxidized to its cognate acid, a structural mimic of auxin [27]. Sirtinol-resistant mutations were found in genes encoding auxin receptors and downstream signalling components as expected, but also in the Moco biosynthesis pathway. This finding indicated that Moco-dependent aldehyde oxidases are likely to catalyze the last oxidation step of auxin. The aldehyde oxidase genes themselves were not found in the mutant screen, most likely because of genetic redundancy of the four *AAO* paralogs [38]. Interestingly, the screen also identified several mutant alleles of *ATM3*, encoding a mitochondrial ABC transporter [23].

In the hope of identifying novel gene products acting upstream or downstream of *ATM3*, we selected two lines, *xd22* and *xd105*, from the pool of uncharacterized sirtinol-resistant mutants based on (i) growth phenotypes found in *atm3* alleles, such as chlorosis and narrow leaves [23], (Figure 2A); (ii) a strong decrease in aldehyde oxidase activity (Figure 2B); and (iii) decreased cytosolic aconitase activity relative to the mitochondrial isozymes (Figure 2C). Ingel activity assays for aldehyde oxidase showed that *xd22* and *xd105* had no detectable activity of the three isozymes that are commonly detected in wild-type leaves (Figure 2B). *xd105* had a strong decrease in cytosolic aconitase activity, concomitant with increased activity of a mitochondrial isozyme. The pattern of relative band intensities for in-gel aconitases in *xd105* was very similar to *atm3-1* (Figure 2C). In the *xd22* line, all three aconitase activities were increased, but the line was taken forward because of its growth phenotype.

**Figure 2.**
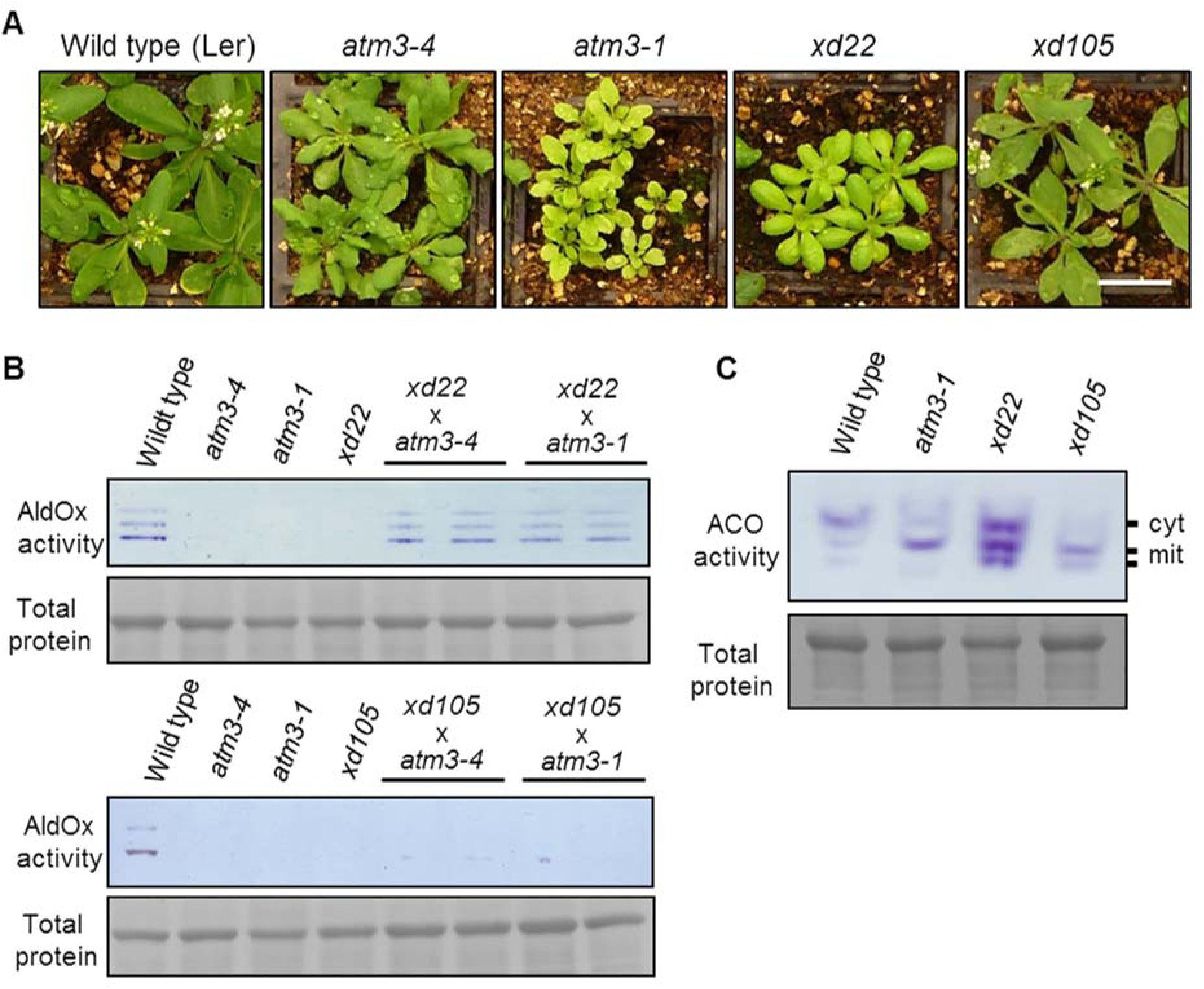
Growth and genetic analysis of *xd22* and *xd105*. A. Four-week-old wild type (Ler) and the indicated mutant lines grown on soil. Scale bar is 2 cm. B. Aldehyde oxidase (AldOx) activities in wild type (Ler), mutant lines and two F1 plants of the indicated crosses. Leaf protein extracts were separated by native PAGE and stained using specific aldehyde substrates and colorimetric electron acceptors. Total proteins were stained to verify equal protein loading. The prominent protein is the large subunit of Rubisco. C. Aconitase activities in wild type (Ler) and mutant lines. Leaf protein extracts were separated by native starch-PAGE and stained for activity using cis-aconitate as substrate, coupled to iso-citrate dehydrogenase and colorimetric electron acceptors. The specific banding pattern corresponding to the cytosolic isoform (cyt, ACO1) and two mitochondrial isoforms (mit, ACO2 and ACO3) was previously reported [17]. Results are representative for three biological repeats (independent plants). Protein loading control as in (B).

To investigate the possibility that the mutation in *xd22* or *xd105* is in the *ATM3* gene, we tested for allelism by crossing with two different *atm3* alleles. *xd22* and *xd105* were used as female parent with *atm3-1* and *atm3-4* as male parent. All mutations are recessive, thus phenotypic rescue is expected if the mutations are not in the same gene. Filial plants were analysed for aldehyde oxidase activities (Figure 2B) and for growth (Figure S1). Aldehyde oxidase activities were fully restored to wild-type levels in the F1 progeny of *xd22* x *atm3* crosses, and the phenotype was comparable to wild type. In contrast, the offspring of *xd105* x *atm3* crosses completely lacked aldehyde oxidase activities, and showed phenotypic similarities to the *xd105* parent with pronounced veins and purple coloration (anthocyanin) at the leaf base (Figure S1). Growth of F1 *xd105* x *atm3-1* seedlings was more vigorous than the parent lines, but this may be due to hybrid effects: *atm3* plants have a defect in DNA repair and accumulate random mutations in their genomes [39], which are outcrossed in the F1. Together, the results suggest that *xd105* contains a deleterious mutation in *ATM3*, whereas the *xd22* phenotype is caused by a mutation in a different gene.

### *xd105* has a mutation in *ATM3* which destabilizes the protein

Sequencing of the *ATM3* gene in *xd105* identified a G1271>A point mutation in exon 13 (Figure 3A). The mutation changes a GGA codon to GAA, leading to substitution of glycine 242 to glutamate (G424>E) in the sixth transmembrane helix (TMH6) of ATM3. The glycine residue is conserved in plant ATM3 and mammalian ABCB7 homologues, but not in *Saccharomyces* or bacterial Atm1 (Figure 3B). To investigate the effect of G424>E on the folding of TMH6, the wild-type and mutant amino acid sequences were run through prediction software for secondary structure. For wild-type ATM3, all six TMH were consistently predicted by four different prediction programs, in agreement with the crystal structure of yeast Atm1 [40]. The G424>E substitution in ATM3 drastically decreased the likelihood of TMH6, see Figure 3C for results from the TMHMM2 server and Figure S2 for results from other servers. To investigate if the mutation in *xd105* affects the stability of ATM3 protein, mitochondria were purified from leaf tissue from wild-type, *xd105* and *atm3-1* plants, and subjected to immunoblot analysis with antibodies against the nucleotide-binding domain of ATM3. The immuno-reactive band corresponding to the 70-kDa monomer of ATM3 could not be detected in the *xd105* line (Figure 3D). The antibodies recognise the N-terminal ATPase domain, including possible ATM3 degradation products shortened at the C-terminus, but no such intermediates were detected. RT-PCR analysis showed that *ATM3* transcript levels in the *xd105* mutant are similar to wild type (Figure 3E). These data suggest that disruption of TMH6 destabilizes the ATM3 protein, possibly by preventing its insertion into the inner mitochondrial membrane. It should be noted that the phenotype of *xd105* is weaker than the *atm3-1* allele lacking the nucleotide binding domain or the knockout line *atm3-2*, suggesting that some functional ATM3 remains which is below the detection limit of the antibodies. Taken together, we conclude that *xd105* is another *atm3* mutant allele and from here on is called *atm3-5*.

**Figure 3.**
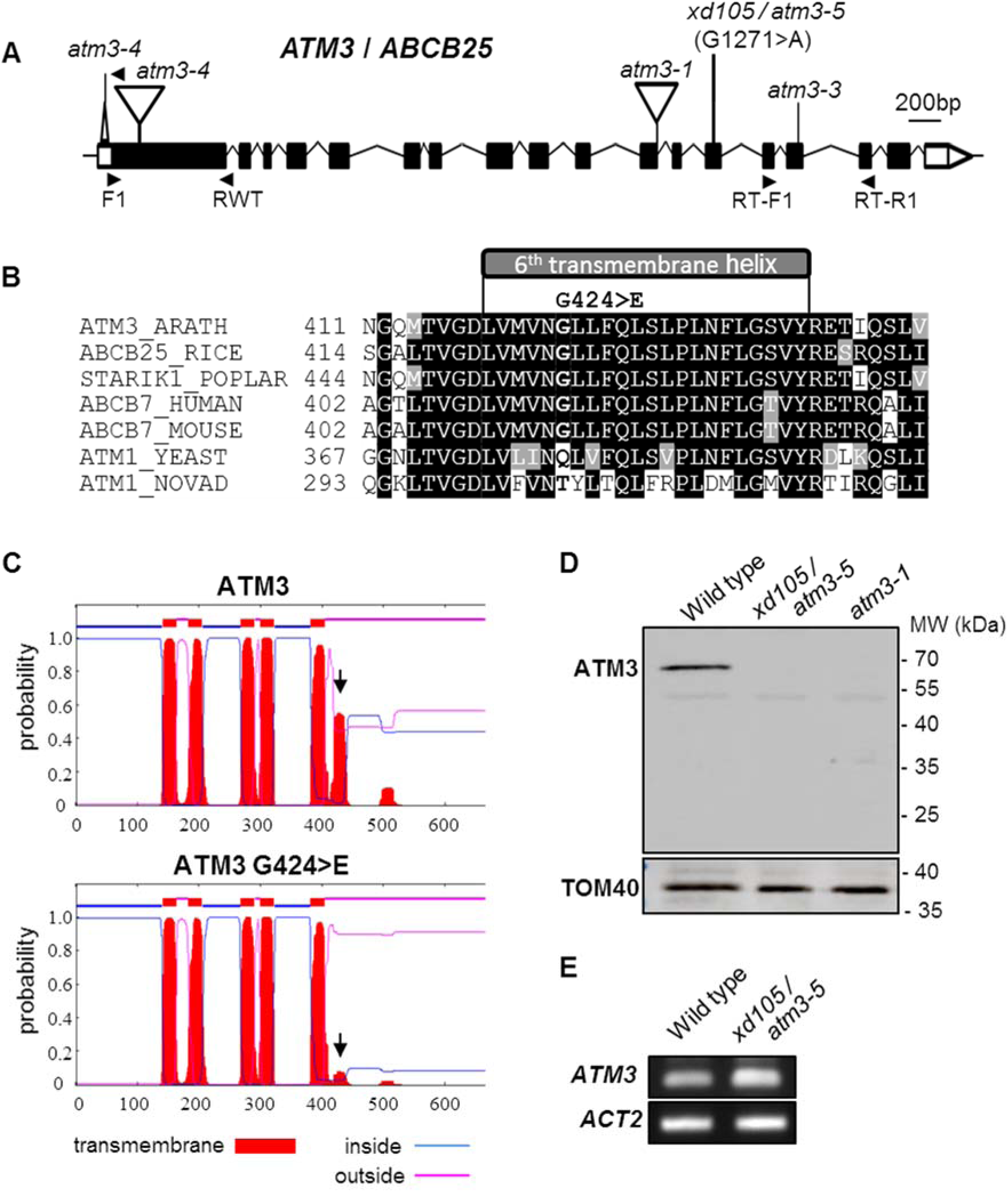
*xd105* has a mutation in *ATM3* leading to destabilization of the protein. A. Gene model of *ATM3* (*ABCB25*, *AT5G58270*) with exons (boxes) and introns (lines). The 3’ and 5’-untranslated regions are in white, coding sequence in black. The G1271>A polymorphism in the *xd105* allele, renamed *atm3-5*, is indicated. B. Amino acid alignment of the sixth transmembrane region of Arabidopsis ATM3. The G1271>A polymorphism is predicted to change glycine 424 into a glutamate (G424>E). Protein sequences were acquired from and aligned in the UniProt database: *Arabidopsis thaliana* (Q9LVM1) (ARATH), *Oryza sativa* subs. *japonica* (Q658I3) (RICE), *Populus balsamifera* subs. *trichocarpa* (B9I784) (POPLAR), human (O75027), mouse (Q61102), *Saccharomyces cerevisiae* (P40416) (SCER), *Novosphingobium aromaticivorans* (Q2G506) (NOVAD). C. Prediction of transmembrane helices in wild-type ATM3 and the G424>E variant using the TMHMM2 server (www.cbs.dtu/services/TMHMM-2.0). The black arrow points to the different probability of transmembrane helix 6. D. Immunoblot analysis of ATM3 in isolated mitochondria from wild-type, *atm3-1* and *xd105 (atm3-5)* seedlings. Immunolabelling with TOM40, the Translocator of the Outer Membrane 40, was used as a control to show equal loading and purity of mitochondria. E. Transcript levels of *ATM3* in 3-week-old wild-type and *xd105* (*atm3-5)* seedlings, using RT-PCR with primers RT-F1 and RT-R1.

### *xd22* has a mutation in *CNX2* affecting the proximal FeS cluster loop

To identify the mutation underlying the *xd22* phenotype, the approximate position of the mutation was determined using SSLP mapping. This showed that *xd22* was located on the right arm of chromosome 2 (Figure S3A). Whole genome sequencing identified a non-synonymous point mutation, G263>A in *CNX2* (Figure 4A, S4B), changing a CGG codon to CAG and causing amino acid substitution of arginine 88 by glutamine (R88Q). R88 is adjacent to a cysteine in the CxxxCxxC motif characteristic of the radical S-adenosylmethionine (SAM) superfamily of proteins. The cysteines coordinate the proximal Fe_4_S_4_ cluster and SAM. Arginine 88 is highly conserved amongst CNX2/MOCS1A/MoaA homologues in plants, fungi, mammals, and bacteria (Figure 4B).

**Figure 4.**
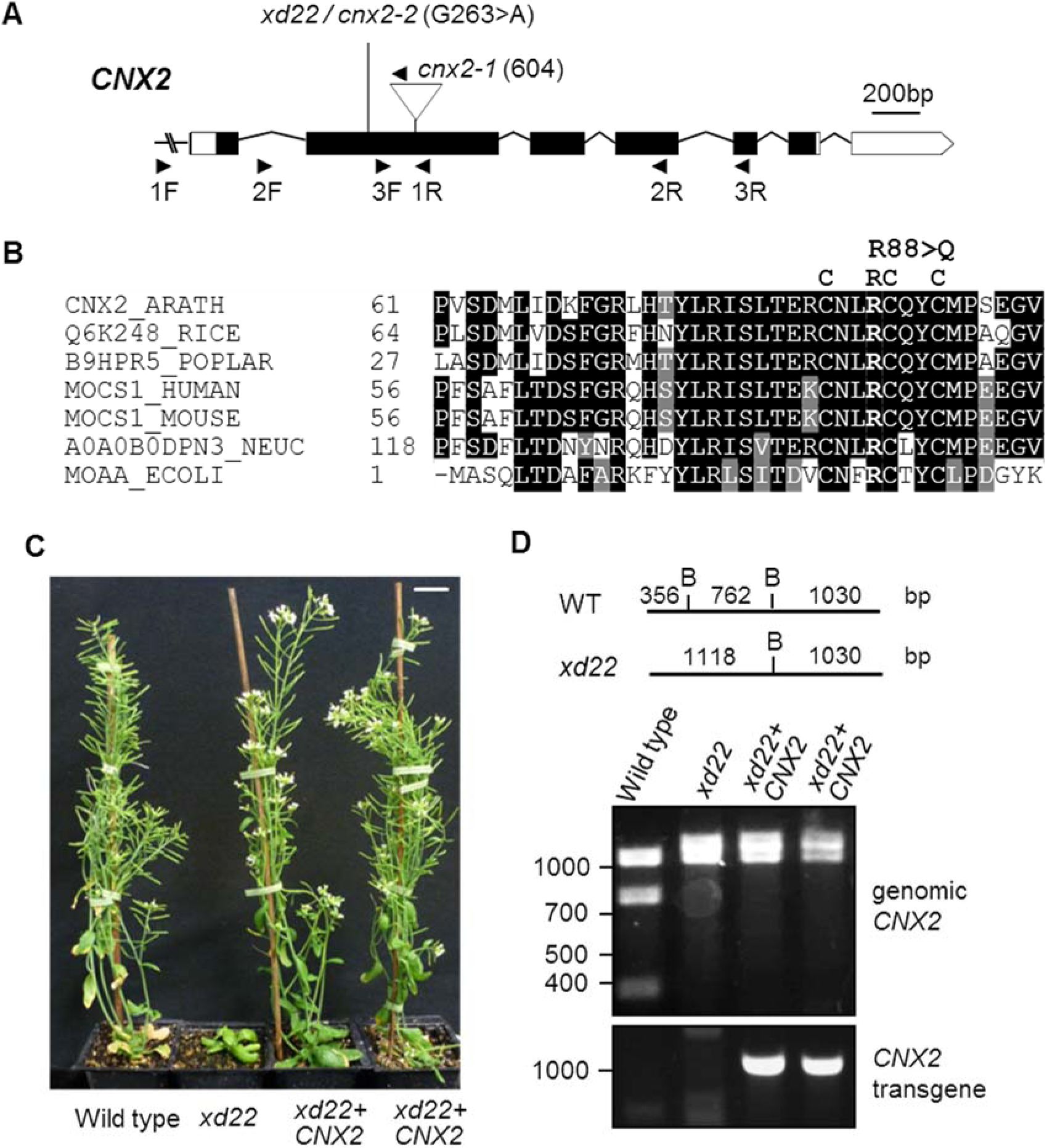
*xd22* has a mutation in *CNX2*. A. Gene model of *CNX2 (AT2G31955)* with exons (boxes) and introns (lines). The 3’ and 5’-UTR are in white, coding sequence in black. The G263>A polymorphism in the *xd22* mutant, renamed *cnx2-2*, is indicated. *cnx2-1* contains a T-DNA insertion (SALK_037143) with the arrowhead indicating the left border primer. B. Amino acid alignment of the N-terminus of Arabidopsis CNX2 and homologs. Protein sequences were acquired from and aligned in the UniProt database: *Arabidopsis thaliana* (Q39055) (ARATH), *Oryza sativa* subs. *japonica* (Q6K248) (RICE), *Populus balsamifera* subs. *trichocarpa* (B9HPR5) (POPLAR), human (Q9NZB8), mouse (Q5RKZ7), *Neurospora crassa* (A0A0B0DPN3) (NEUC), *Escherichia coli* (P30745) (ECOLI). C. Growth of wild-type (WT), *xd22* and two *xd22* plants transformed with *CNX2* (T1 generation), six weeks after sowing. Scale bar is 1 cm. D. Genotype analysis of the plants in (C) to show the presence of the *xd22* polymorphism and the *CNX2* transgene. The 263 G>A mutation in *xd22* removes a BsaWI restriction site (top), altering the restriction pattern of the 2138-nt PCR product obtained with primers CNX2endog 1F and CNX2endog 1R (Figure 4A). The *CNX2* transgene was detected using the primers CNX2trans F and CNX2trans R.

To confirm that the identified polymorphism in *CNX2* is causing the *xd22* phenotype, mutant plants were transformed with a 3-kb genomic sequence containing the wild-type *CNX2* gene including the native promoter. The *CNX2* transgene fully rescued the growth phenotype of *xd22* mutants (Figure 4C, D). To further test if the healthy growth of the complemented line was due to restoration of the Moco biosynthesis defect, aldehyde oxidase activities were measured using the in-gel activity assay. Indeed, plants homozygous for *xd22* containing the wild-type *CNX2* transgene had normal activities of the three main aldehyde oxidase isoforms (Figure 5A). These data prove that *xd22* is a mutant allele of *CNX2* and was renamed *cnx2-2*. To analyse Moco enzyme activities in *cnx2-2*, xanthine dehydrogenase (XDH) and nitrate reductase activities were measured. Xanthine dehydrogenase belongs to the same family of Moco enzymes as aldehyde oxidase, which depend on FAD, two Fe_2_S_2_ cofactors and sulfurated Moco. Using an in-gel enzyme assay with hypoxanthine as substrate, no activity was detectable in leaves from *cnx2-2* plants (Figure 5B). Nitrate reductase is a major plant enzyme depending on haem and Moco and its activity was determined by measuring nitrite production. We found that nitrate reductase activity in leaf extract from *cnx2-2* was 55 ± 5% of wild-type values (Figure 5C). Taken together, genetic complementation and strongly decreased activities of Moco enzymes show that the mutation in *xd22* is located in *CNX2*, encoding the first step in Moco biosynthesis.

**Figure 5.**
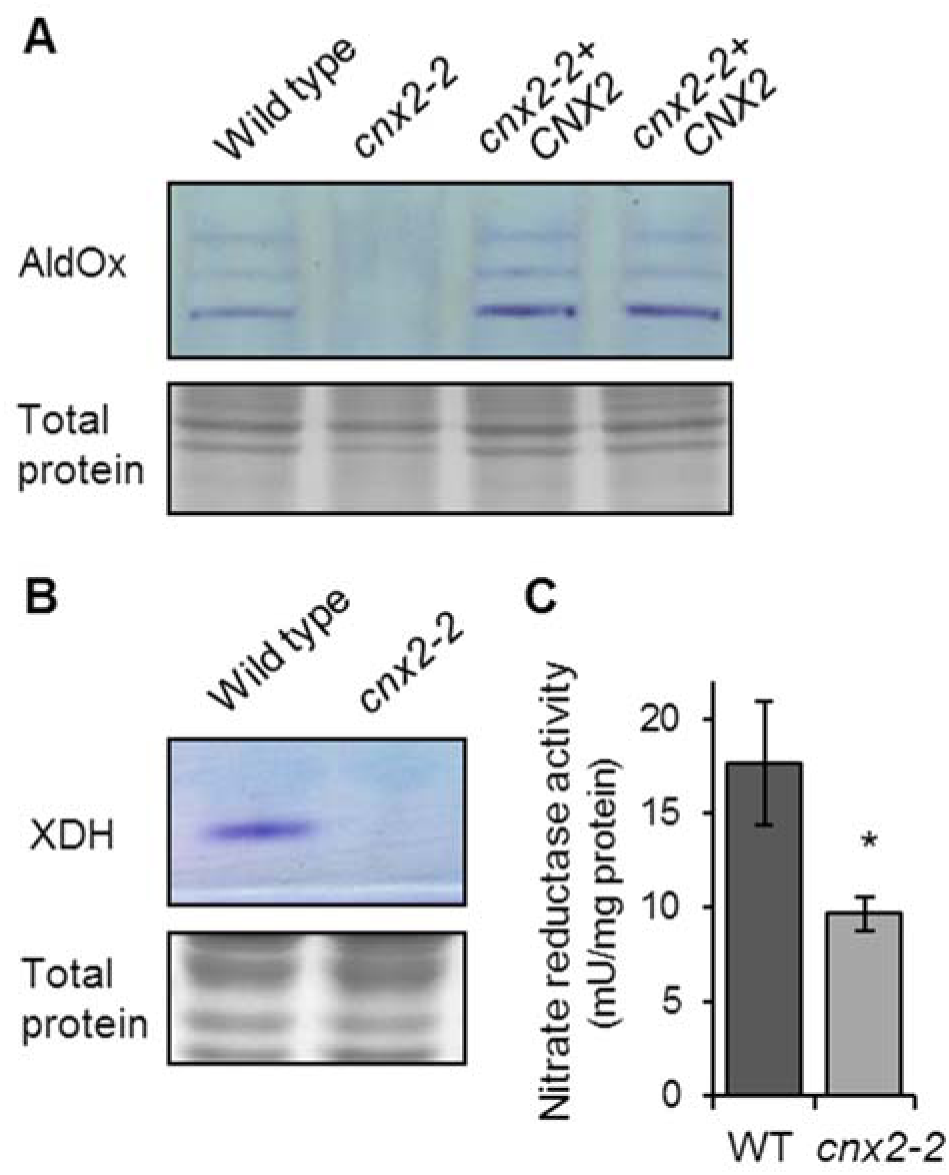
The activities of Moco enzymes are decreased in *cnx2-2* plants. A. In-gel aldehyde oxidase (AldOx) activities in leaves of wild-type, *xd22* and two *xd22* plants complemented with *CNX2* (upper panel). Total protein staining with Coomassie shows equal loading (lower panel). B. In-gel xanthine dehydrogenase (XDH) activity in wild type and *cnx2-2* (upper panel). Total protein staining as in (A). C. Nitrate reductase activity in wildtype (WT) and *cnx2-2* leaves. The values represent the mean ± SE (n = 3), * p > 0.01, Student t-test.

### The *cnx2-2* mutation is not a knock-out allele and growth is rescued by ammonia

To further assess the impact of R88Q substitution on the functionality of CNX2, we compared the *cnx2-2* allele with a knock-out allele. A T-DNA insertion mutant (SALK_037143), from here on named *cnx2-1*, has been reported previously [27], but the line was only minimally characterized. Specifically, there was no genetic confirmation that the T-DNA insertion disrupted the gene, and phenotype analysis was limited to 5-day-old seedlings germinated in the dark. To demonstrate that the T-DNA insertion disrupted expression of *CNX2*, we isolated homozygous *cnx2-1* mutant seedlings from a heterozygous parent (Figure 6A), and performed RT-PCR analysis to confirm that the *CNX2* transcript was absent (Figure 6B). Homozygous *cnx2-1* seedlings segregated in a 1:4 ratio (Figure S4A), which is less than the expected Mendelian ratio of 1:3, but the goodness-of-fit is still significant at p ≤ 0.05 (X^2^ = 4.182).

**Figure 6.**
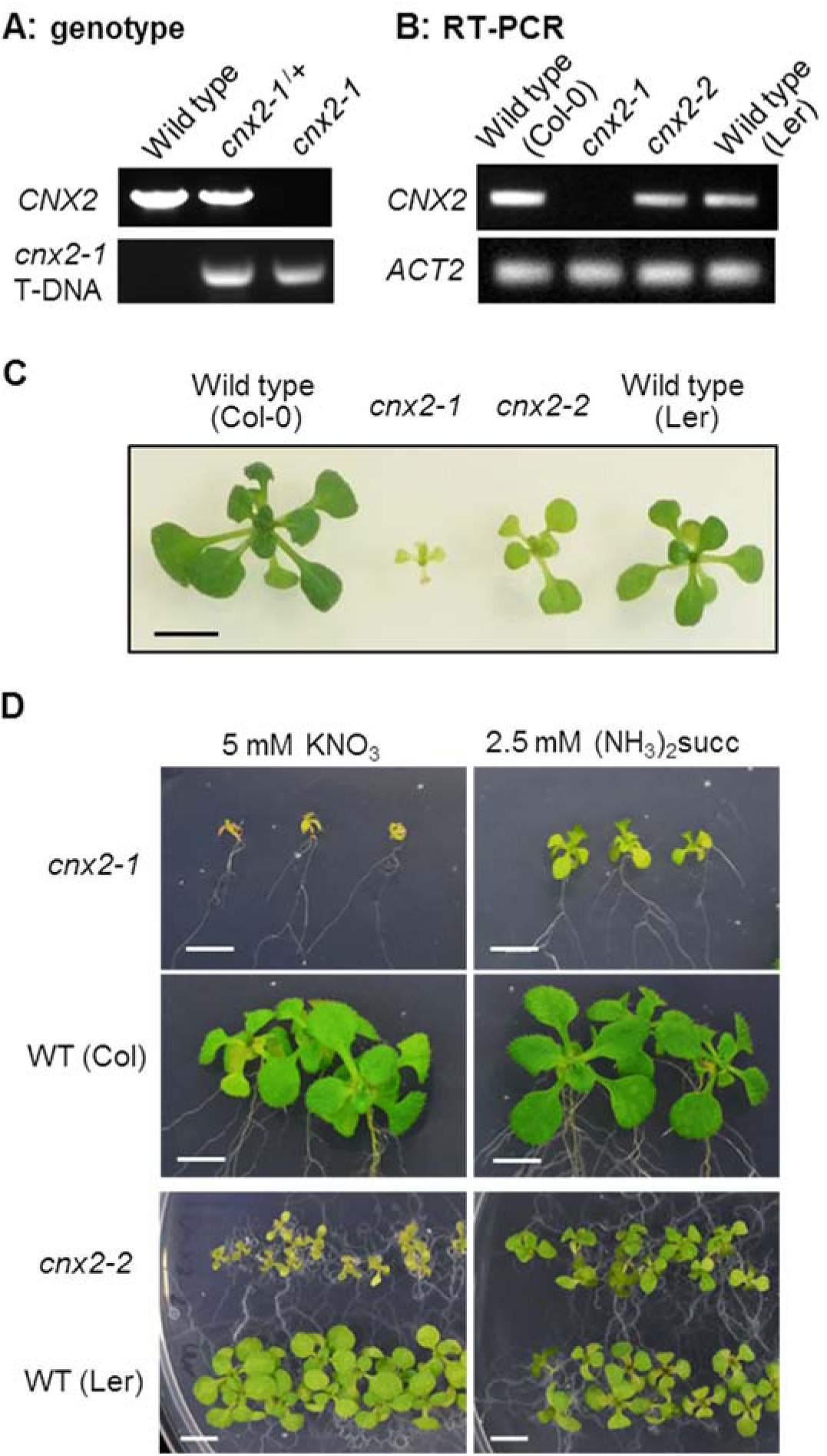
The *cnx2-1* knockout is seedling lethal but growth is rescued by ammonia. A. Genotype analysis of plants in (B). Primer pairs CNX2 2F and CNX2 2R were used to amplify the wild-type product; CNX2 2F and LBb1.3 were used to detect the T-DNA insertion. B. Transcript levels of *CNX2* in wildtype and *cnx2* seedlings, using RT-PCR with primers ACT2 F2 and R2 for *ACTIN2* and CNX2 3F and 3R for *CNX2* amplification. C. Growth of *cnx2-1, cnx2-2* next to their respective wild-type lines on ½MS + 1% (w/v) sucrose for 3 weeks. D. Rescue of growth of *cnx2-1* and *cnx2-2* with ammonium succinate ((NH_3_)_2_succ). Seeds from a heterozygous *cnx2-1*/+ plant were germinated on medium with nitrate (KNO_3_) to select homozygous *cnx2-1* seedlings after 10 days. They were transferred either to fresh medium with NO_3_, or to medium with NH_3_. Images were taken 11 days after transfer. *cnx2-2* seeds and the corresponding wild-type were planted directly on plates with NO_3_ or NH_3_ and images were taken after 14 days. Scale bars in (C) and (D) are 5 mm.

On standard medium with sucrose, the growth of *cnx2-1* seedlings was strongly impaired, the leaves were chlorotic and failed to expand (Figure 6C, S5A). *cnx2-1* seedlings died after two weeks. The phenotype resembles that of plants lacking nitrate reductase [28]. To bypass the requirement of nitrate reductase plants can be grown on medium with ammonium succinate. This nitrogen compound is preferred because, unlike NH4^+^ salts, it does not acidify the medium. When the chlorotic *cnx2-1* seedlings were transferred to medium with ammonia, within a week they recovered chlorophyll levels and started to grow (Figure 6D). *cnx2-2* seedlings germinated directly on ammonia also grew better than on standard medium with nitrate, appearing similar to wild type.

Crosses between heterozygous *cnx2-1 CNX2* and homozygous *cnx2-2* plants gave F1 offspring in two phenotype categories: seedlings with wild-type appearance corresponding to *cnx2-1 CNX2*; and small chlorotic seedlings that were genotyped as *cnx2-1 cnx2-2* (Figure S4B, C). From a single crossing event we obtained 8 wild-type and 11 mutant seedlings, which fits the expected 1:1 ratio although the low numbers are not statistically significant. The *cnx2-1 cnx2-2* seedlings were of intermediate phenotype between *cnx2-2* and *cnx2-1* homozygotes (Figure S4B).

These genetic and phenotypic results show that the R88Q substitution partially inhibits the function of the CNX2 protein, and that lack of nitrate reductase activity is the main growth defect caused by Moco deficiency in Arabidopsis.

### cPMP is decreased in *cnx2* and *atm3*, and accumulates in *cnx5*

To estimate the residual activity of CNX2 R88Q, we set out to measure cPMP, the product of the coupled enzyme activities of CNX2 and CNX3 and their homologs [8]. Following a published method [9], plant extracts were oxidized by KI/I_2_, centrifuged to remove precipitated proteins, and the supernatant was separated by reverse phase HPLC with fluorescence detection. Although we were able to detect synthetic cPMP, it was not possible to resolve cPMP in plant extracts due to its low abundance and overlap with other fluorescent compounds. Therefore, we developed a LC-MS/MS method specific for cPMP, using a HILIC column to get better retention than with a C18 reverse phase column and thus improve separation. The precursor ion *m*/*z* 344 was trapped and 5 different fragments were recorded, of which the *m*/*z* 245.9 transition was most abundant, but *m*/*z* 217.9 was better resolved. Using this method including a calibration range of spiked wild-type samples, we found that Arabidopsis leaf extracts contain approximately 5 nmol cPMP per g extracted protein. A *cnx5* mutant, which is blocked in a downstream biosynthetic step (Figure 1), accumulated 13-fold more cPMP than wild type (Figure 7). In *cnx2-1* and *cnx2-2* mutants cPMP was below the limit of detection, which we estimate to be less than a quarter of wild-type values. We also measured cPMP levels in the *atm3-1* mutant, a relatively strong mutant allele which produces sufficient plant material for analysis (Figure 2A). Interestingly, cPMP levels were approximately 50% decreased compared to wild type. This value correlates well with a previously reported 50% decrease in nitrate reductase activity [9, 23]. Thus, using mass spectrometry rather than fluorescence for the more specific determination of cPMP, we show that cPMP levels are decreased in both *cnx2* and *atm3* mutants.

**Figure 7.**
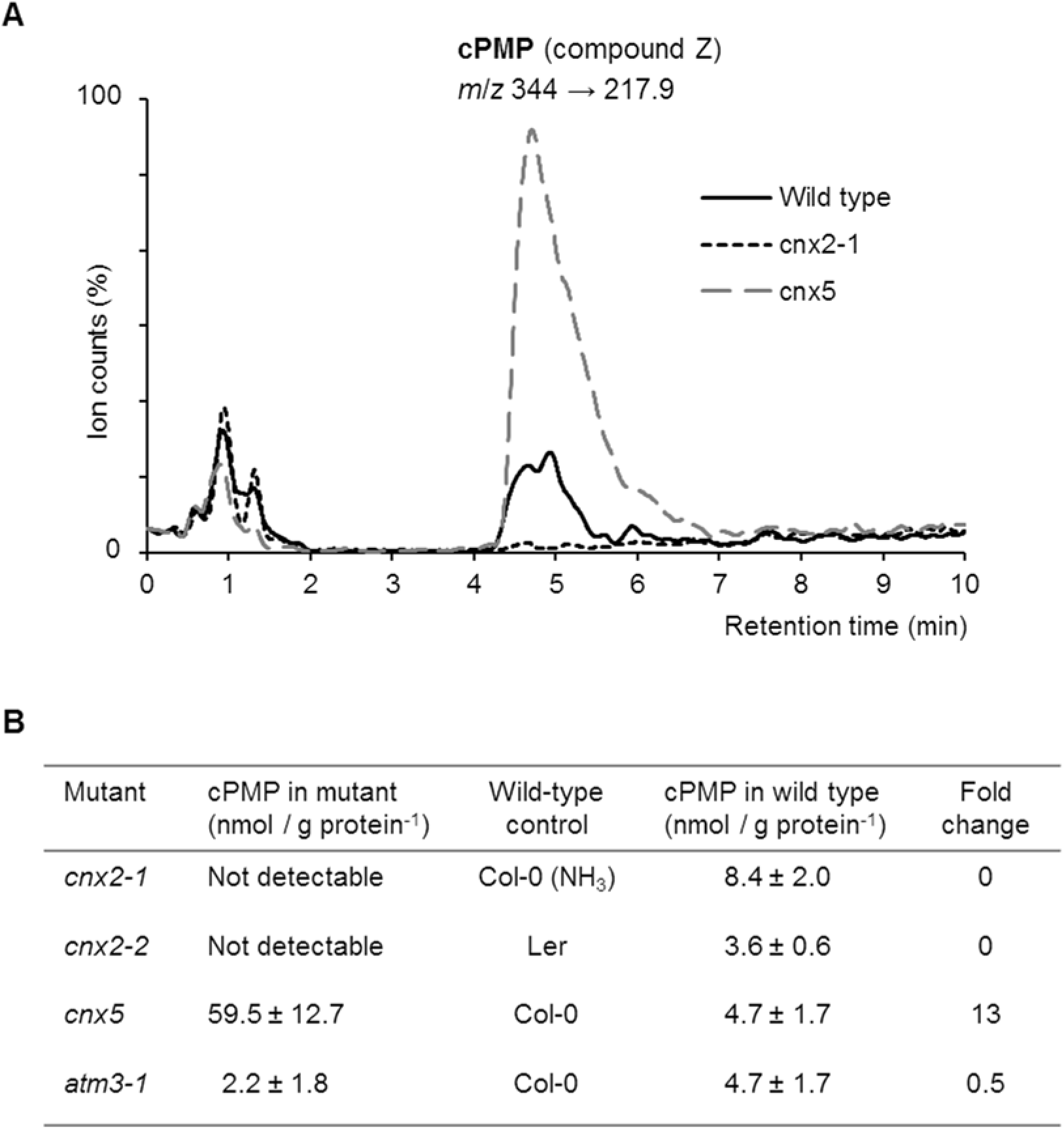
cPMP accumulates in *cnx5*, and is decreased in *cnx2* and *atm3-1* mutants. A. Typical chromatograms of the *m*/*z* 344 to 217.9 transition of cPMP (oxidized to compound Z) in total extracts of wild type (Col-0), *cnx2-1* and *cnx5* seedlings. The protein concentrations of the extracts were 3.7; 4.5 and 5.8 mg / ml, respectively. B. cPMP concentrations in *cnx2*, *cnx5* and *atm3* mutant seedlings, their respective wild-type controls and the fold change compared to wild type. Values represent the mean ± SE (n = 3 – 4 biological replicates of pooled seedlings).

To further compare plant mutants in *ATM3* and *CNX2*, the knock-out alleles *atm3-2* and *cnx2*-1 were grown side-by-side on medium with sucrose but without ammonia. *atm3-2* seedlings are small, but they grew better than *cnx2-1* (Figure S5), indicating that ATM3 is not essentially required for either production of cPMP or a later step in Moco biosynthesis, such as export from mitochondria.

## DISCUSSION

The biosynthesis of Moco has been studied in bacteria, archaea and eukaryotes. In plants, relatively little is known about the first step of the pathway carried out by CNX2 and CNX3, other than their localization in the mitochondrial matrix and a possible involvement of the ATM3 exporter [9]. No targeted mutant studies of *CNX2* and *CNX3* in any plant species have been carried out to our knowledge. We set out to further investigate the function of ATM3 in Moco biosynthesis, which led to identification of a novel *atm3* allele as well as a viable allele of *cnx2*.

The point mutation in the *cnx2-2* allele changes arginine 88 to glutamine in the highly conserved CNLRCQYC sequence that coordinates the proximal FeS cluster using the thiol groups of the 3 cysteines. Crystal structures of the bacterial homolog MoaA [41] show that the side chain of R27, which is equivalent to R88 in Arabidopsis, points away from the FeS cluster, into the loop that positions the cluster in the β-barrel cavity of the enzyme. A glutamine in this position is likely to distort the loop or alter the position of the FeS cluster. This might affect the coordination of S-adenosyl methionine or the distance to the distal FeS cluster. Amino acid changes in the conserved motif have previously been reported to impair the function of the MoaA/MOSC1A protein. Mutating all 3 cysteines to alanines completely abolished enzyme activity [42]. Single amino acid substitutions such as C80G, C84R/F and the adjacent Q85 and Y86D cause Moco deficiency in humans [7,43]. We tried to explore if the R88Q substitution destabilizes the CNX2 protein in Arabidopsis, using protein blot analysis on mitochondrial fractions with previously published antibodies [9]. Unfortunately, we were unable to obtain a specific immuno-reactive signal for wild-type CNX2 with the antiserum, which may have expired over time.

In addition to genetic complementation (Figure 4), we attempted to chemically rescue the *cnx2* mutants with cPMP. In animals, cPMP is injected intravenously and induces a remarkable and sustained improvement of Moco deficiency biomarkers within days after treatment [44]. We used chemically synthesized cPMP [32] and applied this to the roots in the agar medium, or vacuum infiltrated the leaves of *cnx2-2* mutants. Buffers and medium were made oxygen-free to prevent immediate oxidation of cPMP. However, no improvement in growth or diminished chlorosis was seen in the next seven days compared to mock-treated plants. These negative data suggest that cPMP is not taken up by the plant. This fits with our observation that *cnx2-1* seedlings in direct root contact with wild-type seedlings did not grow any better than isolated *cnx2-1* seedlings. Interestingly, *cnx2-1* knockout embryos develop normally in the seed pods of heterozygous mother plants, suggesting that cPMP, MPT or Moco can be transported through vascular cells (symplastic transport). Animal cells deficient in either MOSCS1 or MOSC2 can be co-cultured to rescue each deficiency [24].

In Arabidopsis, mutants in *atm3* are found at a relatively high frequency among Moco biosynthesis mutants in genetic screens for sirtinol resistance ([23]; this study). Sirtinol resistance is caused by decreased activity of aldehyde oxidases, enzymes that depend on FeS and Moco for activity. While several Moco biosynthesis genes have been identified using this genetic screen [27], no genes for FeS cluster assembly have been found thus far, except for *ATM3*. Previous reports showed that stronger mutant alleles of *ATM3* have a decrease in nitrate reductase activity, an enzyme that relies on Moco but not FeS clusters, pointing towards a role for ATM3 in Moco biosynthesis [9,17]. Our LC-MS/MS data showed a decrease in cPMP levels in the *atm3-1* mutant, suggesting that CNX2 or CNX3 activities are affected. Either the expression levels of CNX2 and CNX3 are down, or the enzymes are inhibited. CNX2 has two FeS clusters, but ATM3 is not thought to be required for the assembly of FeS clusters inside the mitochondrial matrix, only for the cytosol [1, 23]. An illustration of this is presented in Figure 2C, showing that the activity of mitochondrial aconitase in the *atm3-1* mutant is normal or even increased, whereas the cytosolic isozyme is inactivated.

However, we know that disruption of ATM3 has an indirect effect on the mitochondrial matrix: using redox-sensitive GFP, we observed that *atm3* mutants have a more oxidized mitochondrial glutathione pool, in agreement with in-vitro transport data that ATM3 exports oxidized glutathione disulfide (GSSG) but not reduced glutathione (GSH) [18]. The chemical reactions carried out by MoaA and MoaC are very sensitive to oxygen [8, 40], and may be less efficient when the redox potential rises to more positive values. In eukaryotes, the homologs of MoaA and MoaC are localized in the mitochondrial matrix, whereas subsequent steps are in the cytosol. Thus, there may be a function reason for cPMP synthesis to take place in the mitochondria, a more reducing environment than the cytosol.

## Abbreviations

ABA3: abscisic acid protein 3
ABC: ATP-binding cassette
ATM3: ABC transporter of the mitochondria 3
AldOx: aldehyde oxidase
cPMP: cyclic pyranopterin monophosphate
CNX: cofactor of nitrate reductase and xanthine dehydrogenase
Col-0: Columbia-0 ecotype
EMS: ethyl methanesulfonate
F1: filial 1
FeS: iron-sulfur
GSSG: glutathione disulfide
GS-S-SG: glutathione trisulfide
ISU/ISCU: iron-sulfur cluster assembly protein 1
Ler: Landsberg ecotype
mARC: mitochondrial amidoxime-reducing component
Moco: molybdenum cofactor
MOCS: molybdenum cofactor synthesis protein
NFS1: nitrogen fixation S (NIFS)-like 1
SAM: S-adenosylmethionine
SNP: single nucleotide polymorphism
SSLP: simple sequence length polymorphism
T-DNA: transfer DNA
TMH: transmembrane helix
TOM40: translocator of the outer membrane 40
URM: ubiquitin-related modifier
XDH: xanthine dehydrogenase

## Acknowledgements

We thank Yunde Zhao (UC San Diego) for making sirtinol-resistant mutants available to the community; Nina Kahlfeldt and Florian Bittner (Technische Universität Braunschweig) for selecting aldehyde oxidase mutants and initial phenotypic characterization; Delphine Bernard for further selecting ‘atm3-like’ mutants; Martin Trick (John Innes Centre) and Zamin Iqbal (EMBL-EBI at Hinxton) for whole genome sequence analysis and SNP identification; Narayan Chaudhuri from Alexion Pharmaceuticals, Inc for providing cPMP; Jim Whelan (La Trobe University) for antibodies against TOM40; Florian Bittner and Christian Gehl for sending antiserum against CNX2.

## AUTHOR CONTRIBUTION STATEMENT

I.K. and A.E.M. designed and performed experiments; L.H. developed the LC-MS/MS method for cPMP and performed the measurements; all authors analysed the data; J.B. wrote the manuscript.

